# A majority of HIV persistence during antiretroviral therapy is due to infected cell proliferation

**DOI:** 10.1101/146977

**Authors:** Daniel B. Reeves, Elizabeth R. Duke, Thor A. Wagner, Sarah E. Palmer, Adam M. Spivak, Joshua T. Schiffer

## Abstract

Antiretroviral therapy (ART) suppresses viral replication in people living with HIV. Yet, infected cells persist for decades on ART and viremia returns if ART is stopped. Persistence has been attributed to viral replication in an ART sanctuary and long-lived and1or proliferating latently infected cells. Using ecological methods and existing data, we infer that >99% of infected cells are members of clonal populations after one year of ART. We reconcile our results with observations from the first months of ART, demonstrating mathematically how a “fossil record” of historic HIV replication permits observed viral evolution even while most new infected cells arise from proliferation. Together, our results imply cellular proliferation generates a majority of infected cells during ART. Therefore, reducing proliferation could decrease the size of the HIV reservoir and help achieve a functional cure.

## Introduction

Antiretroviral therapy (ART) limits HIV replication in previously uninfected cells leading to elimination of most infected CD4+ T cells.^1^ Yet, some infected cells persist and are cleared from the body at an extremely slow rate despite decades of treatment.^2,3^ There is debate whether infection remains due to HIV replication within a small population of cells^4,5^or due to persistence of memory CD4+ T cells with HIV integrated into human chromosomal DNA.^3,6,7^If the latter mechanism predominates, prolonged cellular lifespan and1or frequent cellular proliferation may sustain stable numbers of infected cells.

To optimize HIV cure strategies, mechanisms sustaining infection must be understood. Persistent viral replication in a “sanctuary” where ART levels are inadequate implies a need to improve ART delivery.^8^ If HIV persists without replication as a *latent reservoir* of memory CD4+ T cells, then the survival mechanisms of these cells are ideal therapeutic targets. Infected cell longevity might be addressed by reactivating the lytic HIV replication cycle^9^ and strengthening the anti-HIV cytolytic immune response, leading to premature cellular demise. Anti-proliferative therapies could limit homeostatic or antigen driven proliferation.^10-12^

These competing hypotheses have been studied by analyzing HIV evolutionary dynamics. Due to the high mutation rate of HIV reverse transcriptase and the large viral population size,^13^ HIV replication in the absence of ART produces large viral diversity.^13-15^ Over time, new strains become dominant due to continuous positive immunologic selection pressure against the virus. Repeated “selective sweeps” cause genetic divergence, or a positive molecular evolution rate,^16^ often measured by continual growth in genetic distance between the consensus strain and the founder virus.^17-19^

A recent study documented new HIV mutants during months 0-6 of ART in three participants at a rate equivalent to pre-ART time points. New mutations were noted across multiple anatomic compartments, implying widespread circulation of evolving strains.^4^ One possible explanation for this data is the presence of a drug sanctuary in which ART levels are insufficient to stop new infection events. Alternative proposed interpretations are experimental error related to PCR resampling, or variable cellular age structure within the phylogenetic trees.^20,21^

In other studies of participants on more prolonged ART (at least one year), viral evolution was not observed despite sampling of multiple anatomic compartments.^22-25^Identical HIV DNA sequences were noted in samples obtained years apart,^14,26,27^ suggesting long-lived latently infected cells as a possible mechanism of HIV persistence.^3,6,7,24,25^ Clonal expansions of identical HIV DNA sequences were also observed, demonstrating that cellular proliferation generates new infected cells.^4,12,24,28-3^0 Multiple, equivalent sequences were noted in blood, gut-associated lymphoid tissue (GALT), and lymph nodes, even during the first month of ART.^24,29,30^

The majority of these studies relied on sequencing single genes including *env*, *gag* and *pol*: this approach may overestimate HIV clonality because mutations in other genome segments could go unobserved.^17,31^In addition, these studies also measured total HIV DNA. However, a majority of HIV DNA sequences have incurred deleterious mutations and do not constitute the true replication competent HIV reservoir.^32,33^ To address these issues, a more recent study utilized a comprehensive, whole-genome sequencing approach to confirm the presence of abundant replication competent sequence clones.^34^ In a separate cohort of patients, rebounding HIV sequences arose from replication competent clonal populations.^35^

Another approach to define HIV clonality involves sequencing of the HIV integration site within human chromosomal DNA.^36-40^ While HIV tends to integrate into the same genes,^39,41^ it is extremely unlikely that two cellular infection events would result in HIV integration within precisely the same human chromosomal locus by chance alone.^37^ Thus, integration site analyses abrogate the challenge of overestimating clonality due to incomplete sequencing and provide an elegant surrogate for whole genome sequencing. Previous studies of integration sites found significant numbers of repeated integration sites, providing strong evidence that these infected cells arose from cellular proliferation.^42,43^ These studies are not absolutely conclusive for HIV persistence because integration site sequencing cannot confirm or deny replication competency of the integrated virus.^39^

While HIV sequence clonality has been widely observed, existing studies observed equivalent sequences in a minority (<50%) of observed sequences. Here, we demonstrate that this finding can be explained by incomplete sampling. Using tools adapted from ecology and data from two integration site studies^36,37^and a replication competent HIV DNA study,^34^ we show that nearly all observed unique sequences are likely to be members of clonal populations which derived from cellular proliferation. We predict that the HIV reservoir consists of a small number of massive clones, and a massive number of small clones.

Based on these results, we used a mechanistic mathematical model to reconcile apparent evolution during the early months of ART with apparent clonality after a year or more of ART. The model includes the major proposed mechanisms for HIV persistence: a drug sanctuary, long-lived infected cells, and proliferating infected cells. The model highlights that observed HIV evolution during the first 6 months of ART can be caused by serial observations of long-lived (or proliferated) cells that were once generated by viral replication. We suggest sampling sequences during early ART may result in detection of a positive molecular evolution rate due to the “fossil record” of past infections rather than current viral replication in a drug sanctuary. Based on observed cellular rates, model output after one week of ART shows that a majority of new infected cells are generated by proliferation.

While it remains impossible to rule out a completely unobserved drug sanctuary, our combined approaches suggest that cellular proliferation predominantly drives observed HIV persistence on ART. Consequently, anti-proliferative therapies embody a meaningful therapeutic approach for HIV cure.

## Results

### Defining genetic markers of HIV persistence

During untreated infection, HIV integrates its DNA copy into human chromosomal DNA in each infected CD4+ T cell.^44^ A majority of new infections are marked by novel mutations due to the high error rate of HIV reverse transcriptase and integration into a unique chromosomal location (**Fig 1**). Therefore, continual accrual of new mutations during ART would suggest that ongoing viral replication, perhaps due to inadequate drug delivery to certain micro-anatomic regions, allows HIV to persist during ART.

**Figure 1.**
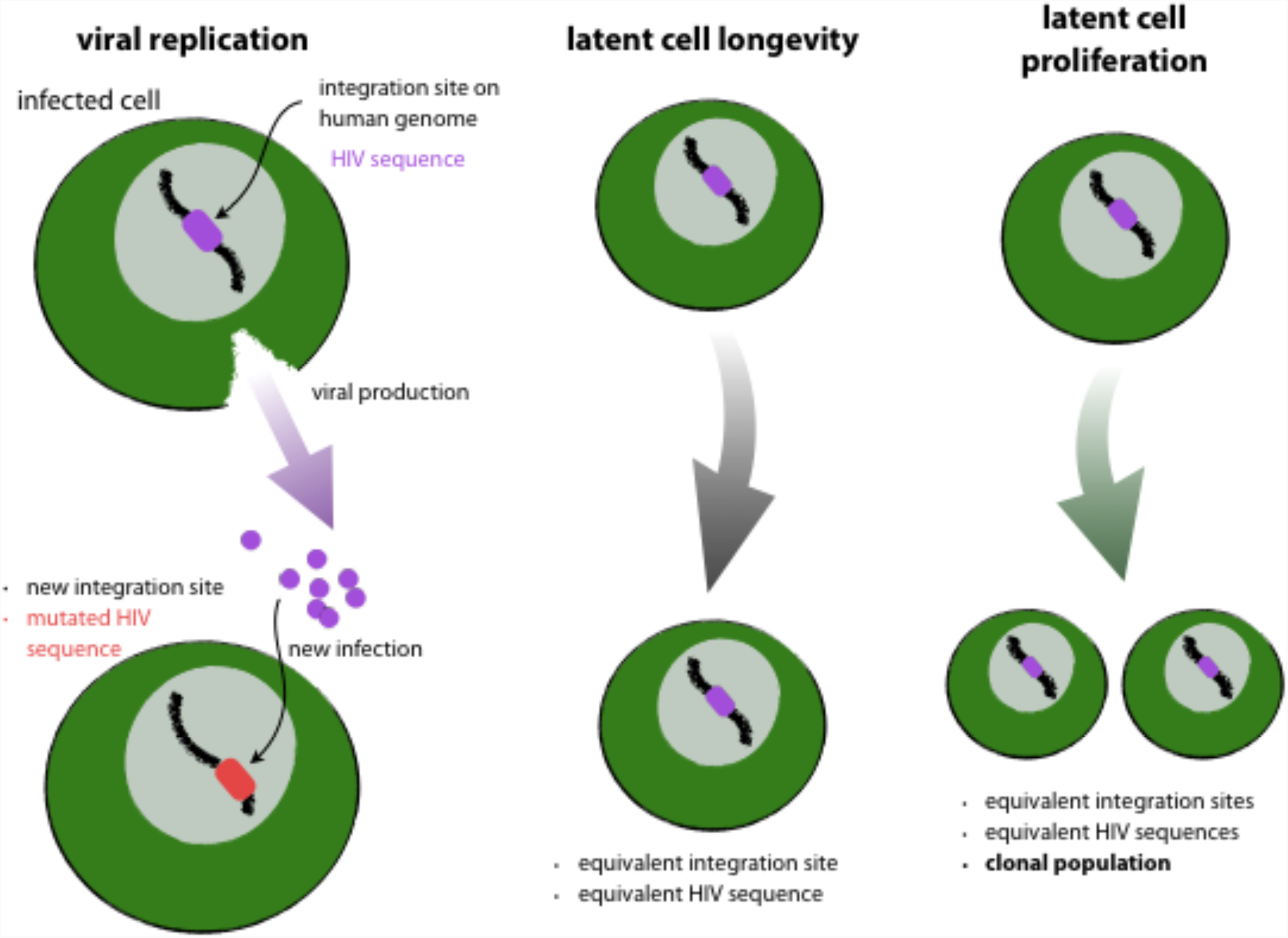
Possible mechanisms for HIV reservoir persistence and their genetic signatures. Viral replication despite ART would lead to accrual of new mutations (color change) and novel chromosomal integration sites in newly infected cells. Alternatively, longevity of latently infected cells maintains sequences and integration sites. Finally, cellular proliferation of latently infected cells produces clonal populations of equivalent HIV sequences and integration sites.

In a subset of infected CD4+ T cells, HIV replication does not progress beyond chromosomal integration and the virus enters latency.^44^ If the same HIV sequences (or integration sites) are found over long time intervals, either cellular longevity or proliferation of latently infected cells allowed HIV to persist. If equivalent HIV sequences with identical chromosomal integration sites are identified in multiple cells, then these viruses were generated via cellular proliferation, rather than HIV replication (**Fig 1**).

Throughout the paper, we contrast the impact of HIV replication and cellular proliferation on HIV persistence during ART by quantifying the numbers or fractions of *unique* sequences and *equivalent* sequences. Human DNA polymerase has much higher copying fidelity than HIV’s reverse transcriptase. Thus, we assume cells whose origin is viral replication will contain unique sequences while cells whose origin is cellular proliferation will contain equivalent sequences and be members of clonal populations.

### Fractions of equivalent total HIV DNA sequences may be extrapolated to replication competent sequences

Most integrated HIV DNA has accrued mutations that render the virus replication incompetent. Quantification of total HIV DNA copies therefore overestimates the size of the replication competent reservoir by 2-3 orders of magnitude relative to viral outgrowth assays.^32^ Replication incompetent, equivalent HIV sequences are commonly present in multiple cells^24,29^. Precisely because these sequences are terminally mutated, they are concrete evidence that some other mechanism (cellular proliferation) copies HIV DNA. The proportion of clonal sequences is similar when analysis includes only replication competent sequences, or all HIV DNA.^34^ As a result, while total HIV DNA may not predict quantity of replication competent viruses, estimates of clonal frequency using total HIV DNA might be extrapolated to the smaller replication competent reservoir.^33^ We use total HIV DNA as it allows a greater sample size for analysis.

### Clonal HIV DNA sequences and clonal replication competent sequences are detectable at various time points during ART

To examine the structure of clonal total and replication competent HIV DNA, we ranked observed sequences from several studies according to their abundance: rank-abundance curves are ordered histograms denoted *a*(*r*) such that *a*(1) is the abundance of the largest clone. These curves facilitate identification of quantities of interest like the richness *R* = max (*r*), sample size *N* = ∑_*.r*_ *a*(*r*), and the number of singletons *N*_1_ = ∑_*.r*_ *I*[*a*(*r*) = 1]. Here *I*[.] is the indicator function equal to 1 when its argument is true and 0 otherwise.

Wagner *et al.* sampled HIV DNA in three participants at three time points 1.1-12.3 years following ART initiation.^37^ Maldarelli *et al.* sampled HIV DNA from five participants at one to three time points 0.2-14.5 years following ART initiation.^36^ In these studies, 1-16% (mean: 7%) of sequences were members of *observed sequence clones* (**Fig 2A**),^36,37^ meaning that HIV DNA was identified in the same chromosomal integration site in at least two cells. The absolute number of observed sequence clones *N*_*i*>1_in the 17 samples ranged from 1-150 (mean: 15). The remaining sequences were identified in a specific chromosomal integration site in only one cell (*observed singletons*).^37^ For total HIV DNA, at each participant time point, certain sequences predominated: the largest observed sequence clone contained 2-62 sequences (mean: 11), accounting for 3-26% (mean= 9%) of total observed sequences.

**Figure 2.**
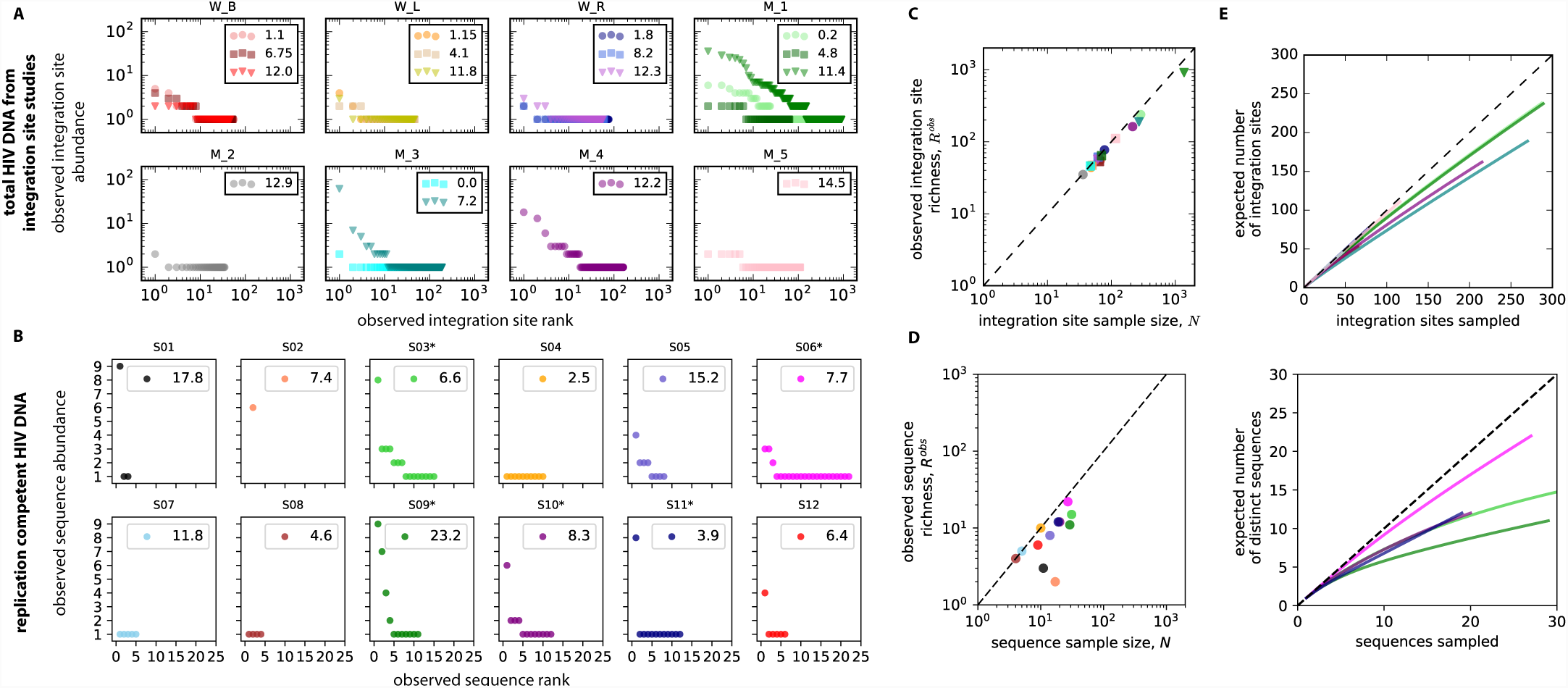
Evidence for clonal HIV sequences. Raw data rearranged as rank abundance curves. **A**. Total HIV DNA from integration site data (Wagner et al., and Maldarelli et al.)^36,37^. Each panel represents a participant, and each marker a duration of ART (indicated in years in the panel legend). W and M in the panel headings distinguish the study. **B**. Replication competent HIV DNA (Hosmane et al.)^34^. Each panel represents a participant. Participants used for analyses below have more than 20 sequences observed (noted by asterisks in panel headings). **C & D.** Sample size of HIV DNA **(C)** and replication competent HIV DNA **(D).** Measuring total HIV DNA increases the number of observed unique sequences (observed sequence richness). The number of total sequences at each time point is plotted against the observed sequence richness. For all HIV DNA samples and when N > 20 for replication competent HIV DNA, the observed richness is always less than the sample size (to the right of the dotted line y=x), owing to the presence of sequence clones. **E.** Sample rarefaction curves for all 17 time points from the 8 study participants in **A** demonstrate the observed number of distinct integration sites as a function of HIV DNA sequence experimental sample size. **F.** Sample rarefaction curves for all 5 study participants in **B** demonstrate the observed number of distinct replication competent HIV DNA sequences as a function of sequence sample size. In both cases, at low sample size, distinct sequences are commonly observed with each new sample. As sample size increases, distinct sequences are increasingly less likely to be detected owing to the presence of repeatedly detected sequence clones. As more and more unique sequences are detected, the curves would flatten until all unique sequences are detected and the curve is completely flat.

Hosmane *et al.* sequenced replication competent HIV isolates from 12 study participants on ART: 0-28% (mean: 11%) of sequences were members of *observed sequence clones* (**Fig 2B**).^34^ The lack of detected clones in 3 participants may reflect their low sequence sample size. Participants with fewer than 20 total sequences were therefore excluded from individual analyses described below but were included for population level evaluations. For replication competent HIV DNA in the 5 persons having sequence sample-size *N* > 20, certain sequences dominated: the largest observed sequence clone contained 3-9 sequences (mean: 6.8), accounting for 11-42% (mean= 28%) of total observed sequences. The number of non-singleton sequence clones *N*_*i* >1_in the 5 samples ranged from 1-7 (mean: 3.8).

### Sequence sampling depth is low relative to total population size

There was a higher number of experimentally detected sequences (*N*) for total HIV DNA (**Fig 2C**) than for replication competent HIV (**Fig 2D**). For total HIV DNA, the number of observed *unique* sequences (*R*^*obs*^or the *observed sequence richness*) was always less than *N* (**Fig 2C**) due to clonal populations. Where *N* > 20 for replication competent viruses, *R*^*obs*^ was always less than *N*, again due to the presence of clones **(Fig 2D)**. There was a higher *R*^*obs*^ as the sequence sample size increased (**Fig 2C&D**), suggesting that detection of unique clones increases with deeper sampling.

Thus, we can infer that further sampling would likely uncover new unique sequences. To quantify the relationship between sample size and discovery, we generated sample rarefaction curves (see **Methods** and **Supplementary Methods**) using the rank-abundance distributions (**Fig 2E&F**). These curves interpolate the data to demonstrate the likely discovery of new sequences as sampling increases up to the sample size of the original experiment. At low sample size, a new sequence is likely to be found with each additional sample. As sampling increases, the chance of sampling a previously documented sequence increases, and the slope of the rarefaction curve begins to flatten. As sample size approaches the true richness of the population, the curve plateaus and few new unique sequences remain to be sampled. Current sampling depth remains on the steep, initial portion of the curve.

### Ecological estimates of lower bounds on true HIV sequence richness from limited samples

To estimate a lower bound for true sequence richness, we used the Chao1 estimator, a nonparametric ecologic tool that uses frequency ratios of observed singletons *N*_1_and doubletons *N*_2_ (see **Methods** and **Supplementary Methods**).^45,46^For the HIV reservoir, theoretical values for true richness range from one (if all sequences were identical and originated from a single proliferative cell) to the total population size (if all sequences were distinct and originated from error-prone viral replication). We found estimated lower bounds for true sequence richness exceeded observed richness, typically by an order of magnitude in both total HIV DNA and replication competent HIV (**Fig 3**). These initial lower bound estimates for sequence richness are far lower than previously estimated population sizes for HIV DNA and replication competent HIV DNA sequences,^2,3,6^ suggesting that clones may predominate.

**Figure 3.**
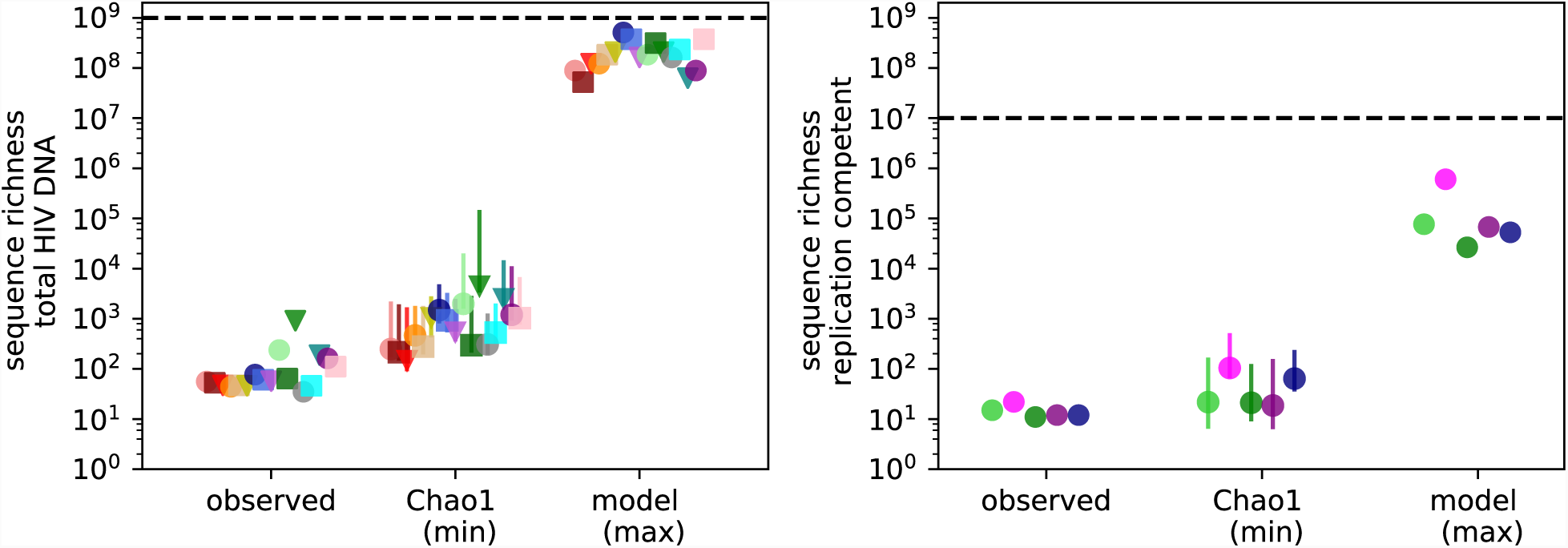
The actual total number of distinct HIV sequences far exceeds the observed total number of distinct HIV sequences during ART. Observed sequence richness underestimates the true HIV sequence richness. For both data sources, Chao1 provides an estimate of the lower bound (min) of true sequence richness (error bars are asymmetric confidence intervals, see **Supplementary Methods**). In all cases, Chao1 estimates are above observed values. Our modeling technique estimates a much higher upper bound (max) for true sequence richness. Nevertheless, the total HIV sequence population size (dashed lines: 10^9^for total HIV DNA and 10^7^ for replication competent HIV) is 1-2 orders of magnitude above the upper bound estimates for sequence richness, suggesting substantial clonality of HIV sequences.

### A majority of observed HIV sequences are members of large proliferative clones

The Chao1 estimator does not include information about the total population size. However, estimates for the total number of total DNA and replication competent sequences in the entire reservoir exist.^33^ Using that additional information, we developed an ecologic model to extrapolate the true rank-abundance of HIV sequences for each participant time point.

Based on the observation that observed data was roughly log-log-linear **(Figure 2A)**, we chose a power-law model for rank-abundance: *a*(*r*) ∝r^*-α*^Other functional forms were explored (exponential, linear, and biphasic power law) but were worse or equivalent for data fitting (not shown). Our model requires 3 parameters, the power law exponent (*α*), the sequence population size (*L*), and the sequence richness (*R*). Model fitting is described in the **Methods** with additional detail in the **Supplementary Methods**. Briefly, we generated 2,500 possible models for each data set, choosing a plausible fixed population size from available data (*L* = 10^9^ for HIV DNA and *L* = 10^7^ for intact, replication competent HIV DNA).^2,3,6,33,47^ We then recapitulated the experiment by taking N random samples from each model distribution and comparing sampled data to experimental data to find optimal model parameters. This resampling method correctly inferred the power law exponent from simulated power law data (**Supplementary Fig 1**).

However, for experimental data we could not precisely identify *R*. Recognizing this uncertainty, we developed an integral approximation to estimate the largest possible richness (least clonality) given *L* and the best-fit *α* (derivation in **Supplementary Methods** and illustration in **Supplementary Fig 2**). Then, using the lower bound estimate from the Chao1 estimator, we were able to fully constrain the estimate of true HIV sequence richness in the reservoir. Our maximal estimates for sequence richness were notably several orders of magnitudes higher than Chao1 estimates (**Fig 3**) but lower than the total sequence population size (*L*).

Our method demonstrated excellent fit to cumulative proportional abundances of observed clones for total HIV DNA (**Fig 4A**) and replication competent HIV DNA (**Fig 5A**). For total HIV DNA (**Fig 4B**) and replication competent HIV DNA (**Fig 5B**), optimal fit was noted within narrow ranges for the power law slope parameter but across a wide possible range of true sequence richness. Using the top 5 best fit models, we generated extrapolated distributions of the entire HIV sequence rank-abundance for each participant time point. We observed similar estimates for the population size of the largest clones, which account for approximately 50% of the reservoir (200-2,000 clones for HIV DNA in **Fig 4C** and 2-7 clones for replication competent HIV DNA in **Fig 5C**). However, the tail of the reservoir, which consists of thousands of smaller clones, varied considerably across the parameter sets with 900-100,000 possible clones accounting for 90% of the HIV DNA and 100-2,000 possible clones accounting for 90% of replication competent HIV. This variability reflects the fact that true sequence richness is only partially identifiable using our procedure.

**Figure 4.**
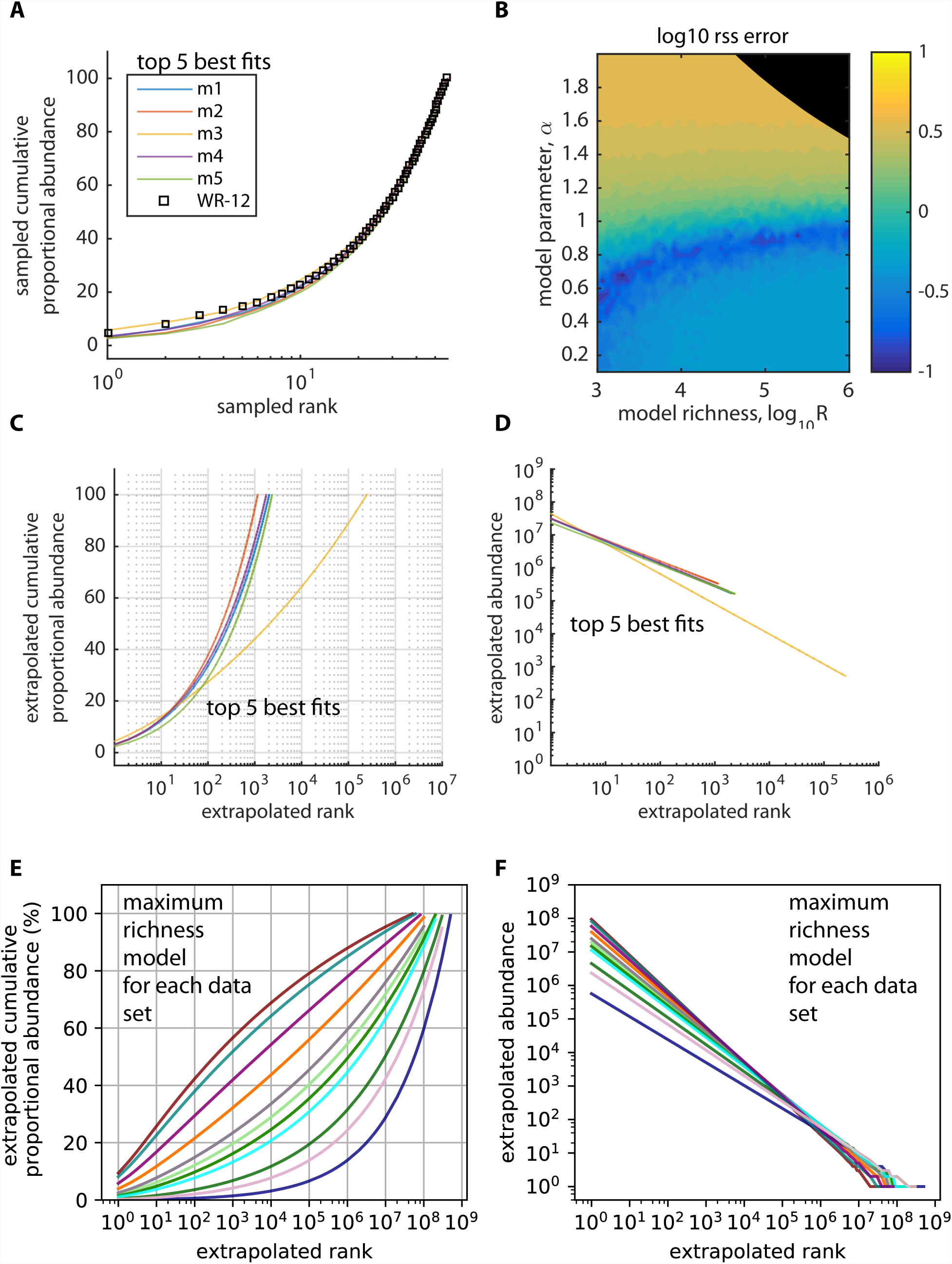
Ecologic modeling suggests a majority of HIV DNA sequences are members of sequence clones. To model the true rank abundance distribution of the HIV reservoir, we used a power law model and recapitulated experimental sampling (sample size equal to the experimental sample size) from 2,500 theoretical power law distributions to fit the best model to participant data in **Fig 2A**. Theoretical distributions varied according to the slope of the power law and the true sequence richness but were fixed at 10^9^ total HIV DNA sequences. **A.** Five best model fits to cumulative proportional abundance curves from a single representative participant (WR, 12 years on ART). Black circles represent the experimental data; the 5 colored model lines are superimposed based on virtually equivalent fit to the data. **B.** Heat diagram representing model fit according to power law exponent α and true sequence richness R with best fit noted by minimum error score (blue color, see details of calculation in results above); black shaded areas represent parameter sets excluded based on the Chao1 estimator (lower bound on sequence richness) and mathematical constraints of the power law (upper bound for sequence richness). A wide range of values for sequence richness allow excellent model fit while the power law exponent is well defined. **C.** Extrapolations of the best-fit cumulative distribution function to the entire pool of 10^9^ infected cells; under the most conservative estimates, the top 200,000 ranked clones constitute the entire reservoir. **D.** Extrapolations of the best fit power law to the entire pool of 10^9^ infected cells; the top 1000 clones consist of >10^4^ cells each. **E.** Extrapolations of the best fit cumulative distribution function to the entire pool of 10^9^ infected cells for all participant time points in **Fig 2A**; we assume the maximum possible sequence richness in each case and still note a predominance of sequence clones. **F.** Extrapolations of the best-fit power law to the entire pool of 10^9^ infected cell for all participants in **Fig 2A;** the top 1,000 clones each consist of >10^4^ cells each. A large number of clones (~10^6^) contain many fewer cells (<100).

**Figure 5.**
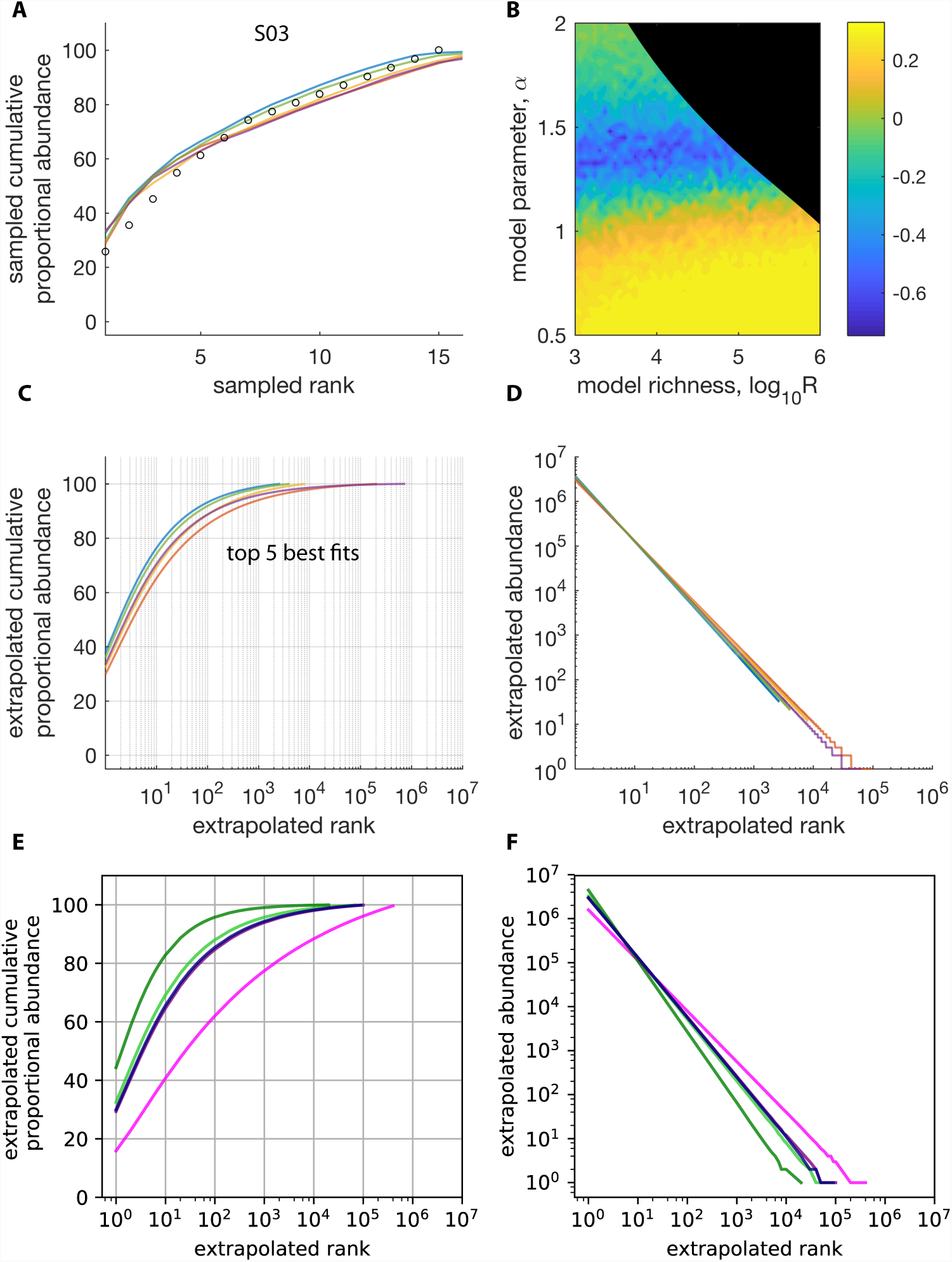
Ecologic modeling suggests a majority of replication competent HIV sequences are members of sequence clones. To recapitulate experimental conditions in **Fig 2B**, we performed in silico sampling (sample size equal to the experimental sample size) from 2,500 theoretical power law distributions of replication competent HIV clone size distributions sorted by rank. Theoretical distributions varied according to the exponent of the power law model and the true sequence richness and were fixed at a reservoir size of 10^7^ replication competent HIV DNA sequences. **A.** Five best model fits to cumulative proportional abundance curves from a single representative participant (S10). Black circles represent the experimental data; the 5 colored model lines are from five separate parameter sets. **B.** Heat map representing model fit according to power law slope α and true sequence richness R with best fit noted by lowest error (blue color); the black shaded area represents parameter sets excluded based on mathematical constraints of the power law (upper bound on sequence richness). A wide range of values for sequence richness (<10^5^ sequences) allow excellent model fit while power law slope falls within a narrow range. **C.** Extrapolations of the best-fit cumulative distribution function to the entire pool of 10^7^ infected cells; under the most conservative estimates, the top 10^5^ ranked clones constitute the entire reservoir. **D.** Extrapolations of the best fit power law to the entire pool of 10^7^ infected cells; the top 100 clones consist of >10^4^ cells each. **E.** Extrapolations of the best fit cumulative distribution function to the entire pool of 10^7^ infected cells for all participants and time points (see original data in **Fig 2B**); we assume the largest possible observed sequence richness in each case and still note a predominance of sequence clones. **F.** Extrapolations of the best-fit power law to the entire pool of 10^7^ infected cell for all participants in **Fig 2B;** the top 100 clones again consist of >10^4^ cells each. A large number of clones (~10^4^) contain many fewer cells (<100).

Even under the most conservative assumptions (maximum possible true sequence richness in **Fig 3**), the vast majority of sequences were predicted to be members of true sequence clones. For the participant in **Fig 4C**, a maximum of 200,000 clones were needed to reach 100% cumulative abundance for HIV DNA. The ratio of estimated true sequence richness to the total number of infected cells R1L with HIV DNA (~10^5^: 10^9^) represents an upper limit on the fraction of sequences that are true singletons: we estimate that greater than 99.9% of infected cells contain true clonal sequences (**Fig 3**).

Similarly, the ratio of estimated true sequence richness to the total number of infected cells with replication competent HIV for the participant in **Fig 5C** was 10^5^:10^7^. Hence, at least 99% of cells contain true clonal sequences (**Fig 3**). Of note, this ratio is stable regardless of assumed reservoir size. For instance, if we assume a true reservoir size of 10^6^, then our estimate of true sequence richness is ~10^4^.

The model fitting procedure was used on all data in **Fig 2**. We biased against a clonally dominated reservoir to the greatest extent possible by selecting the best fitting power law exponent and then calculating the maximum possible sequence richness (**Fig 3**). The power law slope parameter was on average lower across participants for HIV DNA (*α* = 0.9 ± 0.1) than for replication competent HIV DNA (*α* = 1.4 ± 0.2). As a result, the predicted cumulative distribution of HIV DNA (**Fig 4E**) was often concave-up with log rank as compared to concave-down with log rank noted for replication competent HIV DNA (**Fig 5E**), suggesting that a smaller number of extremely large clones might make up a higher proportion of the replication competent HIV reservoir.

For both HIV DNA (**Fig 4F**) and replication competent virus (**Fig 5F**), the top 100 clones in all participants are estimated to be massive (>10^5^ and >10^4^ cells respectively). However, there are also large numbers of much smaller clones with fewer than 1,000 cells (>10^6^ and >10^4^ clones respectively). In contrast to observed data, a majority of sequences are clonal, suggesting that proliferation is the major generative mechanism of persistent HIV-infected cells.

### Modeling combined population data gives similar results as individual fitting

To increase sample size and eliminate bias related to excluding participants with low sample sizes, we combined results from all participant time points for HIV DNA (17 time points) and replication competent HIV (12 time points) into single rank order distribution curves. We then fit the power law models to both sets of data (**Supplementary Fig 3A&B, E&F**). We again noted a narrow range of possible values for the power law exponent and a large range of possible values for true sequence richness. The exponent was again *α* < 1 for total HIV DNA and *α* ≈ 1 for replication competent virus (**Supplementary Fig 3A&E**), leading to concave-up and linear relationships between cumulative proportional abundance and log rank, respectively (**Supplementary Fig 3C&G**). We estimated that at least 99.9% of cells with HIV DNA (**Supplementary Fig 3C**) and 99.8% of cells with replication competent HIV (**Supplementary Fig 3G**) contain true clonal sequences. The top 100 HIV DNA clones (**Supplementary Fig 3D**) and replication competent clones (**Supplementary Fig 3H**) contained >10^6^ and >10^4^ cells respectively.

Using the population level data, we generated sample rarefaction curves from the extrapolated rank-abundance curves. These curves show that after 10,000 sequences were sampled, the observed sequence richness would continue to increase with more sampling (**Supplementary Fig 4**). Even if experimental sample sizes could be increased 100-fold from the present data, sequences would continue to be dominated by those from large clones. Our statistical inference approach is therefore necessary to provide a more realistic estimate of the clonal distribution of the HIV reservoir.

### A mechanistic model that includes both an ART sanctuary and cellular proliferation can reconcile observations from early and late ART

Our analyses above identify the critical role of cellular proliferation in generating infected cells after a year of ART but do not capture the dynamic mechanisms underlying this observation or explain possible evidence of viral evolution during months 0-6 of ART.^4^ We therefore developed a viral dynamic mathematical model. Our model (**Fig 6A**) consists of differential equations, described in detail in the **Methods**. Most model parameter values are obtained from the literature (**Table 1**).

**Table 1:** Model parameters

| Parameter | Value | Meaning | Units | Source |
| --- | --- | --- | --- | --- |
| $R_0$ | 8 | Basic reproductive number of HIV | [] | 69 |
| $\beta_0$ | $2 \times 10^{-4}$ | Viral infectivity, used in $\beta_\epsilon = \beta_0(1 - \epsilon)$ | $[\mu\text{L copy}^{-1} \text{day}^{-1}]$ | 61,70,71 |
| $\epsilon$ | 0.95 | ART efficacy outside the sanctuary | [] | 71,72 |
| $\pi$ | $10^3$ | Viral production rate, used in $n = \pi/\gamma$ | $[\mu\text{L copy}^{-1} \text{day}^{-1}]$ | 61,70,71 |
| $\gamma$ | 23 | Viral clearance rate, used in $n = \pi/\gamma$ | $[\text{day}^{-1}]$ | 73 |
| $\alpha_S$ | 150 | Susceptible cell production rate | $[\mu\text{L copy}^{-1} \text{day}^{-1}]$ | 54,70,71 |
| $\delta_S$ | 0.2 | Susceptible cell death rate | $[\text{day}^{-1}]$ | 61,70,71 |
| $\delta_1$ | 0.8 | Productively infected cell ($I_1$) clearance rate | $[\text{day}^{-1}]$ | 71,74 |
| $\delta_2$ | 0.02 | Pre-integration cell ($I_2$) death rate | $[\text{day}^{-1}]$ | 48,50 |
| $\alpha_2$ | 0.047 | Pre-integration cell proliferation rate | $[\text{day}^{-1}]$ | 42 |
| $\xi_2$ | 0.08 | Pre-integration cell activation rate | $[\text{day}^{-1}]$ | Fit |
| $\alpha_{3(j)}$ | [0.047,0.015, 0.002] | Proliferation rate of latently infected cells $j \in [T_{em}, T_{cm}, T_n]$ phenotypes, respectively | $[\text{day}^{-1}]$ | 42 |
| $\xi_3$ | 0.0003 | Latent cell activation rate (for all $j$) | $[\text{day}^{-1}]$ | Fit |
| $\delta_{3(j)}$ | | Calculated from latent clearance rate as $\theta_L = \alpha_{3(j)} - \delta_{3(j)} - \xi_3$ where $\theta_L = -5.2 \times 10^{-4}$ | $[\text{day}^{-1}]$ | 2,3 |
| $\tau_{i(j)}$ | [1, $10^{-2}$, $10^{-4} \varrho_j$] | Probability of infection of each compartment, taken from y-intercepts in Ref. 50 | [] | 51 |
| $\varrho_j$ | [0.2,0.75,0.05] | Fraction of latent infected cells of each phenotype (e.g. from patient #5 in Ref. 12) | $[\text{cells } \mu\text{L}^{-1}]$ | 12 |
| $V(0)$ | $10^2$ | Initial viral load (from typical set-point value $10^5$ copies/mL) | $[\text{copy } \mu\text{L}^{-1}]$ | 69 |
| $I_1(0)$ | 2 | Initial concentration of productively infected cells, calculated from $I_1(0) = V(0)/n$ | $[\text{cells } \mu\text{L}^{-1}]$ | 75 |
| $I_2(0)$ | 0.2 | Initial concentration of pre-integration infected cells | $[\text{cells } \mu\text{L}^{-1}]$ | 75 |
| $I_{3(j)}(0)$ | $0.2\varrho_j$ | Initial concentration of each latent phenotype, calculated from $\sim 10^6$ latently infected cells in $\sim 5\text{L}$ of blood | $[\text{cells } \mu\text{L}^{-1}]$ | 2,12 |
| $I_S(0)$ | 180 | Initial concentration of sanctuary cells, calculated from equilibrium model $I_S(0) = \frac{\alpha_S}{\delta_1} - \frac{\delta_S}{n\beta_0(1-\epsilon_S)}$, e.g. Ref. 56 SI | $[\text{cells } \mu\text{L}^{-1}]$ | Calc |
| $\zeta$ | 0.007 | Decay rate of T cell activation | $[\text{day}^{-1}]$ | 52 |
| $\epsilon_S$ | 0 | ART efficacy in the sanctuary, minimum value | [] | Min |
| $\varphi_S$ | $10^{-5}$ | Fraction of cells in sanctuary | [] | 4 |

**Figure 6.**
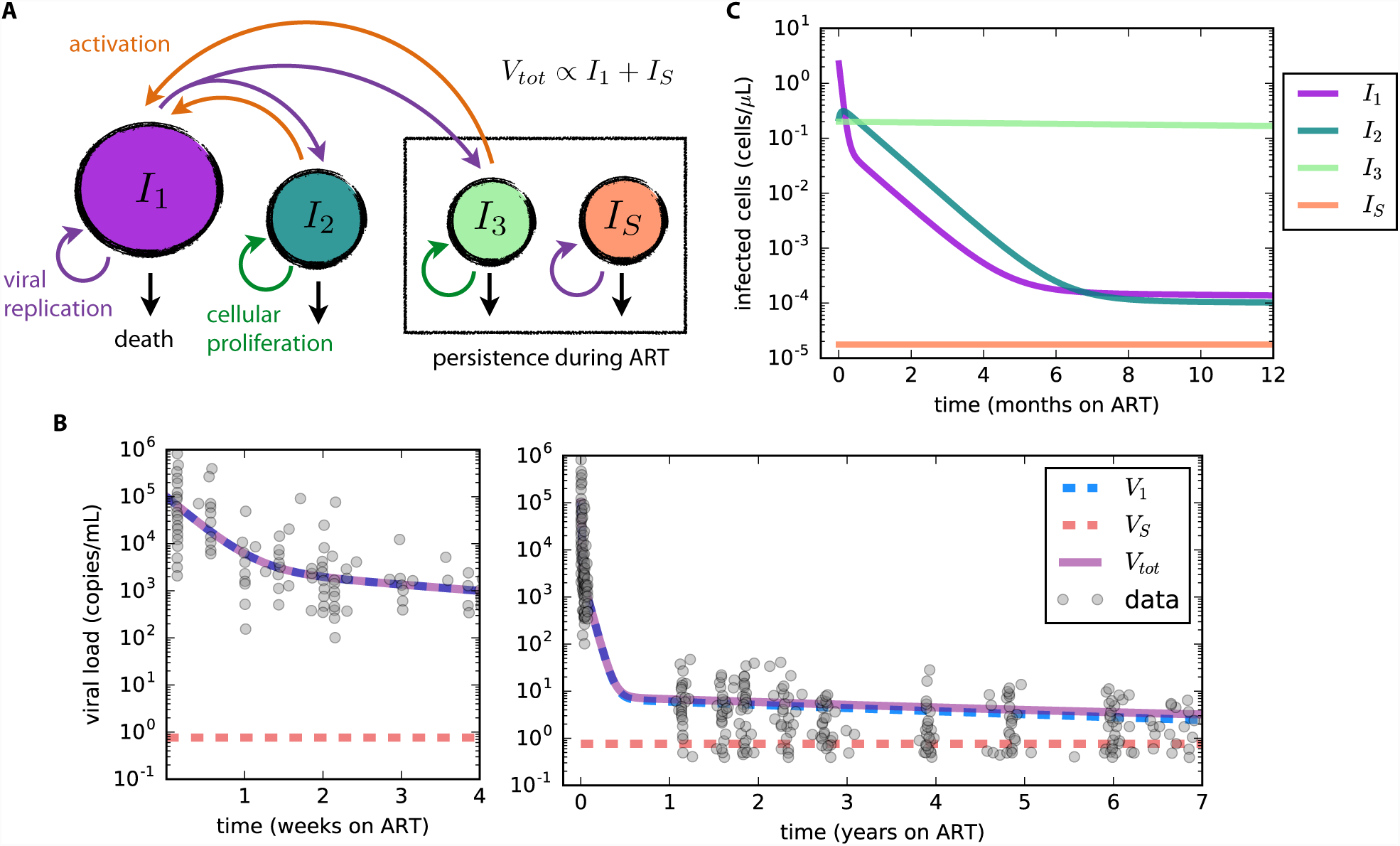
A mechanistic model recapitulates HIV RNA decay and predicts rough equivalence of virus produced by the sanctuary and virus produced by a reactivating reservoir up until months 4-6 of ART. **A**. Model schematic: I_1_cells produce virus, pre-integration latent cells I_2_are longer lived and eventually transition to I_1_ and long-lived latently infected cells I_3(j)_proliferate and die at measured rates depending on cell phenotype j (e.g. effector memory, central memory, naive. Sanctuary cells I_s_ allow ongoing HIV replication despite ART. Parameters and their values are discussed in the Methods and listed in Sup Table 1. **B**. The mathematical model recapitulates observed HIV RNA data (Palmer et al.^51^) over weeks and years of ART. V_1_ is virus derived from I_1_ while V_s_ is derived from I_s_. **C**. I_2;_ and I_3_ become the predominant cell types early during ART. I_s_ remains very low throughout the duration of ART which is necessary to explain the lack of detectable viremia on fully suppressive ART.

Briefly, we classify rapid death *δ*_1_ and viral production within actively infected cells *I*_1_. Cells with longer half-life *I*_*2*;_ are activated to *I*_1_ at rate *ξ* _2_. *I*_2_ may represent CD4+ T cells with a prolonged pre-integration phase, but their precise biology does not affect model outcomes.^48^ The state *I*_3 (*j*)_ represents latently infected reservoir cells of phenotype *j*, which contain a single chromosomally integrated HIV DNA provirus.^44^ *I*_*3*_ reactivates to *I*_1_ at rate *ξ*_3_.^49^ The probabilities of a newly infected cell entering *I*_1_, *I*_2_ *I*_*3* (*j*)_, are *τ*_1_, *τ*_2_*τ*_3 (*j*)_. Because we are focused on the role of proliferation, we assume sub-populations of *I*_*3*_,^12^ including effector memory (T_em_), central memory (T_cm_), and naïve (T_n_) CD4+ T cells, which have been experimentally proven to turn over at different rates _3 (*j*)_, *δ*_3 (*j*)_.^12,42,43^

ART potency *є* ∈ [0,1] characterizes decrease in viral infectivity due to ART.^50^ Other dynamic features of infection such as death rate of infected cells, latent cell proliferation rate and reactivation rates of latent cells, are unchanged on ART. In our simulations, the basic reproductive number becomes *R*_0_ (1 -*є*) on ART and is <1 when *є* > 0.95, meaning that each cell infects fewer than one other cell and viral load declines from its previous steady state until becoming undetectable. Only short stochastic chains of new infection can occur.

To make a model inclusive of viral evolution despite ART, we allow for the possibility of a drug sanctuary state (*I*_*s*_) that reproduces with reproductive number *R*_0_ (1 -*є*_*s*_)~8. In the drug sanctuary, ART potency is assumed to be negligible (*є*_*s*_= 0) such that the sanctuary reproductive number is equivalent to the value from a model without ART. Target cell limitation or a local immune response must result in a sanctuary viral set point to prevent infected cells and viral load from growing exponentially. The sanctuary size must also be limited (0.001-0.01% of the original burden of replicating HIV) to achieve realistic viral decay kinetics.^51^ In the absence of contradictory information, we assumed homogeneous mixing of *V*_1_ and *V*_*s*_ in blood and lymph nodes.^4^

Based on the observation that activated, uninfected CD4+ T cells (*S*), the targets for replicating HIV, decrease in numbers after initiation of ART we also simulate the model with and without the possibility of slow target cell decline within the HIV drug sanctuary. We approximate this process with an exponential decay of target cells with rate *ζ*(per day).^52,53^ The decay rate is lower than concurrent decay rates measured from HIV RNA^50,51,54^ because abnormal T cell activation persists for more than a year after ART.^53^

### The model accurately simulates viral dynamics during ART

We fit the model to ultra-sensitive viral load measurements collected from multiple participants in Palmer *et al*.^51^ We included experimentally derived values for most parameter values (**Table 1**), solving only for activation rates *ξ*_2_and *ξ*_3_by fitting to viral load. Simulations reproduce three phases of viral clearance (**Fig 6B**) and predict trajectories of infected cell compartments (**Fig 6C**). Of note, the model is able to achieve fit to the data with different assumptions of starting values of the three infected cell compartments (the relative proportion of which are unknown pre-ART): in this circumstance, we arrive at different values of *ξ*_2_and *ξ*_3_ without impacting overall model conclusions regarding the HIV reservoir. The size of the sanctuary (expressed as the fraction of infected cells *φ*_*s*_) is only constrained to be below a value <10^-5^ to ensure accurate model fit for a static sanctuary model.

### Cellular proliferation sustains HIV infection during ART whether or not a small drug sanctuary exists

We next used the model to estimate the fraction of cells generated by cellular proliferation versus viral replication. We conservatively assumed that prior to ART all infected cells were generated by viral replication. Then, we tracked the number of cells whose origin was replication and the number whose origin was cellular proliferation. Without directly simulating a phylogeny, the fraction of all cells that derive from replication provides a surrogate for the expected fraction of cells that would give a signal of evolution. We also distinguish the *current replication percentage*, the fraction of infected cells currently being generated from viral replication, from the *net replication percentage*, the fraction of total infected CD4+ T cells at a given time whose origin was HIV replication. This distinction allows us to contrast the net number of surviving, historically-infected cells with the number of cells that are presently being generated via HIV infection. Because many long-lived cells were once generated by HIV infection, the net replication percentage may exceed the current replication percentage.

We then simulated the model under several plausible sanctuary and reservoir conditions to assess the relative contributions of infection and cellular proliferation in sustaining infected cells. We considered different reservoir compositions based on evidence that effector memory (T_em_), central memory (T_cm_) and naïve (T_n_) cells proliferate at different rates and that distributions of infection in these cells differ among infected patients.^12,42,43^ Further, because a drug sanctuary has not been observed, its true volume is unknown and may vary across persons. We therefore conducted simulations with a static sanctuary, a slowly diminishing sanctuary, and no drug sanctuary **(Fig 7A)**.

**Figure 7.**
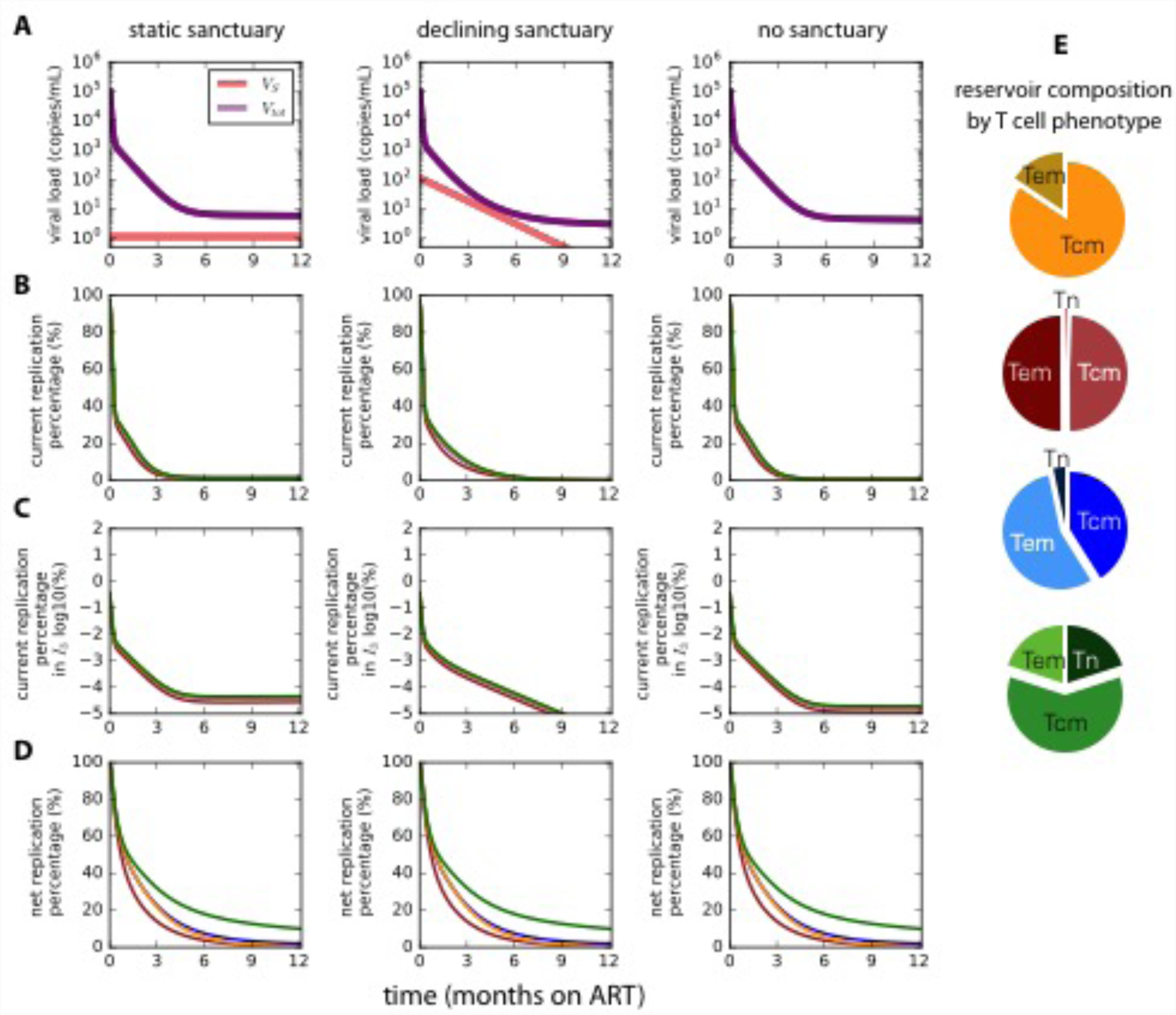
The vast majority of infected cells are generated via proliferation within 6 months of ART initiation. Model simulations contrast the number of cells generated by viral replication with those generated by cellular proliferation. The fraction of cells generated by replication at any time point is referred to as the current replication percentage. The fraction of cells that remain alive whose ultimate origin was viral replication is referred to as the net replication percentage. Different assumptions regarding sanctuary (I_s_) and latent cell populations (I_3_) were simulated corresponding to columns. **A.** Moving left to right, we assume a static drug sanctuary, a slowly declining drug sanctuary and no drug sanctuary. Pie charts on the right indicate the reservoir composition by T cell phenotypes and correspond with colored lines in **B-D. B.** Under all assumptions, once ART is initiated, most new infected cells arise due to cellular proliferation as opposed to HIV replication after 12 months of ART. **C.** New latently infected reservoir cells (I_3_) are generated almost entirely by proliferation soon after ART is initiated under all conditions. **D.** The observed proportion of infected cells originally generated by HIV infection rather than cellular proliferation will overestimate the actual ongoing proportion during the first 6 months of ART assuming a small or large sanctuary volume. This trend is more notable when the reservoir contains a higher proportion of slowly proliferating naïve T cells.

Regardless of assumed pre-treatment reservoir composition and sanctuary size, the contribution of replication to generation of new infected cells is negligible after one year of ART. The contribution of new replication diminishes rapidly with time on ART regardless of whether a sanctuary is assumed (**Fig 7B**). The fraction of long lived latently infected cells (*I*_*3*_) generated by viral replication (**Fig 7C**, note log scale) is negligible within days of ART initiation. This finding captures the extent of the impact of proliferation even when a sanctuary is assumed.

### Observable HIV DNA sequence evolution during early ART can represent a fossil record of prior replication events

In all simulations, the net fraction of cells generated from viral replication rather than cellular proliferation at 6 months of ART (5-25% in **Fig 7D**) is higher than the current percentage generated by replication (**Fig 7B**). A higher fraction of slowly proliferating T_n_ cells exacerbates the difference between historical and contemporaneous generation of infected cells (**Fig 7D**, green line). Because the net fraction is what will be observed experimentally, the model reveals why ongoing evolution might be observed even while the dominant mechanism sustaining the reservoir is cellular proliferation. In keeping with the first section of our paper, after 12 months of ART, the net and current percentage of infected cells generated by HIV replication become negligible for all simulated parameter sets. Importantly, the lag between net and current viral replication generation emerges whether or not a small drug sanctuary is included in the model.

We refer to the phenomenon that long-lived cells may contain signatures of past viral replication as the “fossil record”. To emphasize the concept, the fossil record finding is qualitatively illustrated in **Fig 8** using a population of 30 infected cells. At 3 time points following the initiation of ART, we compare the net and current percentage of cells generated by viral replication. At day 60, 30% of cells remain that were originally generated by viral replication. This means 30% of observed sequences might produce a signal of evolution. However, at that time an overwhelming majority of new infected cells are being generated by proliferation.

**Figure 8.**
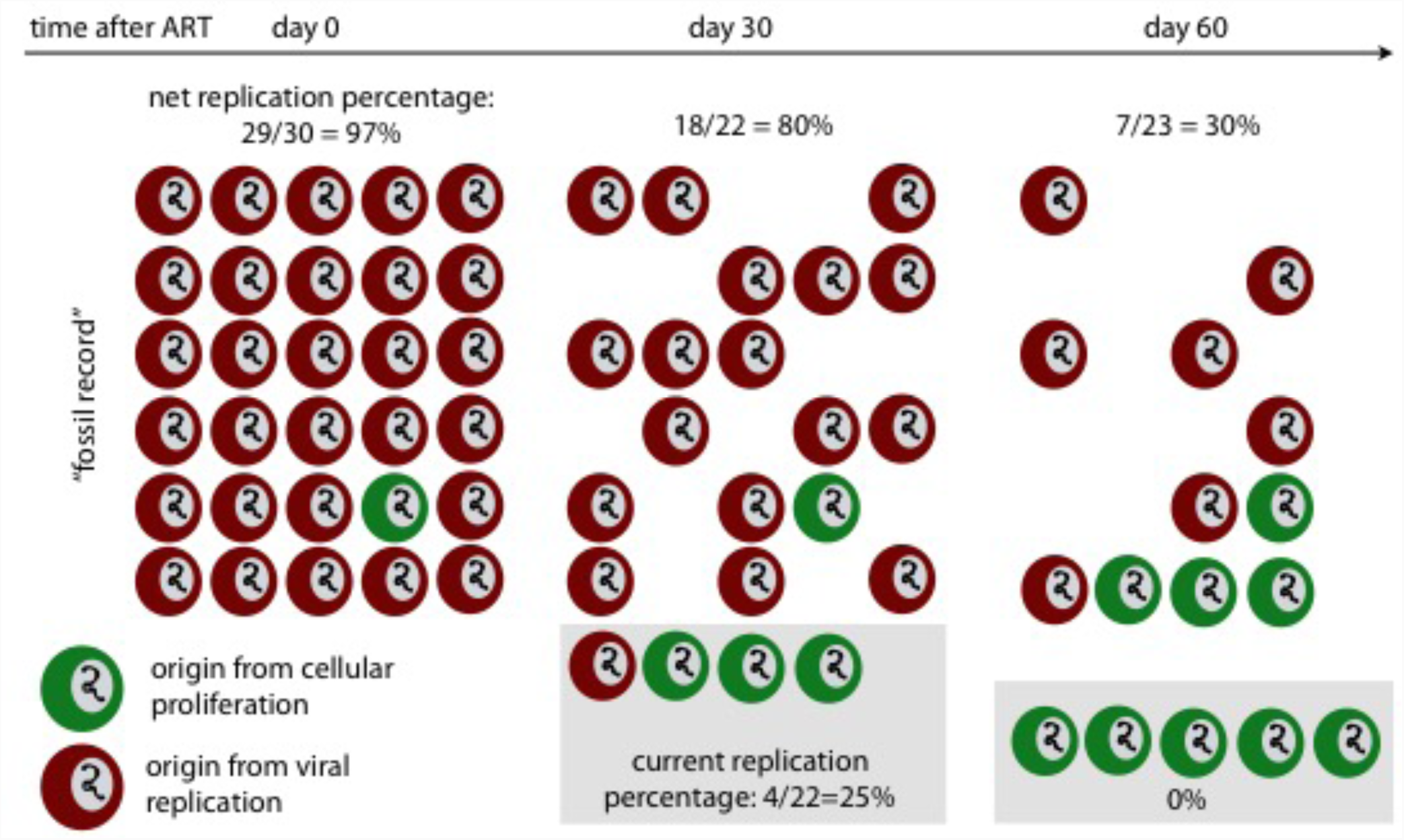
Qualitative illustration of the fossil record phenomenon. In an example population of 30 infected cells, the proportion of infected cells that were once generated by HIV replication (the net replication percentage, or “fossil record” of HIV replication) remains >30% for the first 2 months of ART. However, in this time, the proportion of cells newly generated by HIV replication (shaded box) becomes negligible. The net fraction is observed experimentally, so our simulations indicate a contemporaneous representation of the HIV reservoir cannot be observed until the “fossil record” is completely washed out, sometime between 6 months and a year of ART.

### Different factors drive net (observed) and current replication percentage during early ART

We next performed sensitivity analyses to identify parameters that impact the timing of transition from HIV replication to cellular proliferation as a source for new and observed infected cells. Under all parameter assumptions, the majority of new infected cells arose from proliferation after a year of chronic ART (**Fig 9A**). Only the sanctuary decay rate (*ζ*) had an important impact on generation of new infected cells. Our analysis included a sanctuary in which target cell availability did not decay at all. In that scenario, 5-10% of new infected cells were generated by HIV replication after a year of ART (**Fig 9A**), which is not consistent with lack of viral evolution observed at this timepoint. Rapid disappearance of HIV replication as a source of new infected cells was identified regardless of initial reservoir volume, drug sanctuary volume, ART efficacy, and reservoir composition (fraction of T_em_, T_cm_, and T_n_).

**Figure 9.**
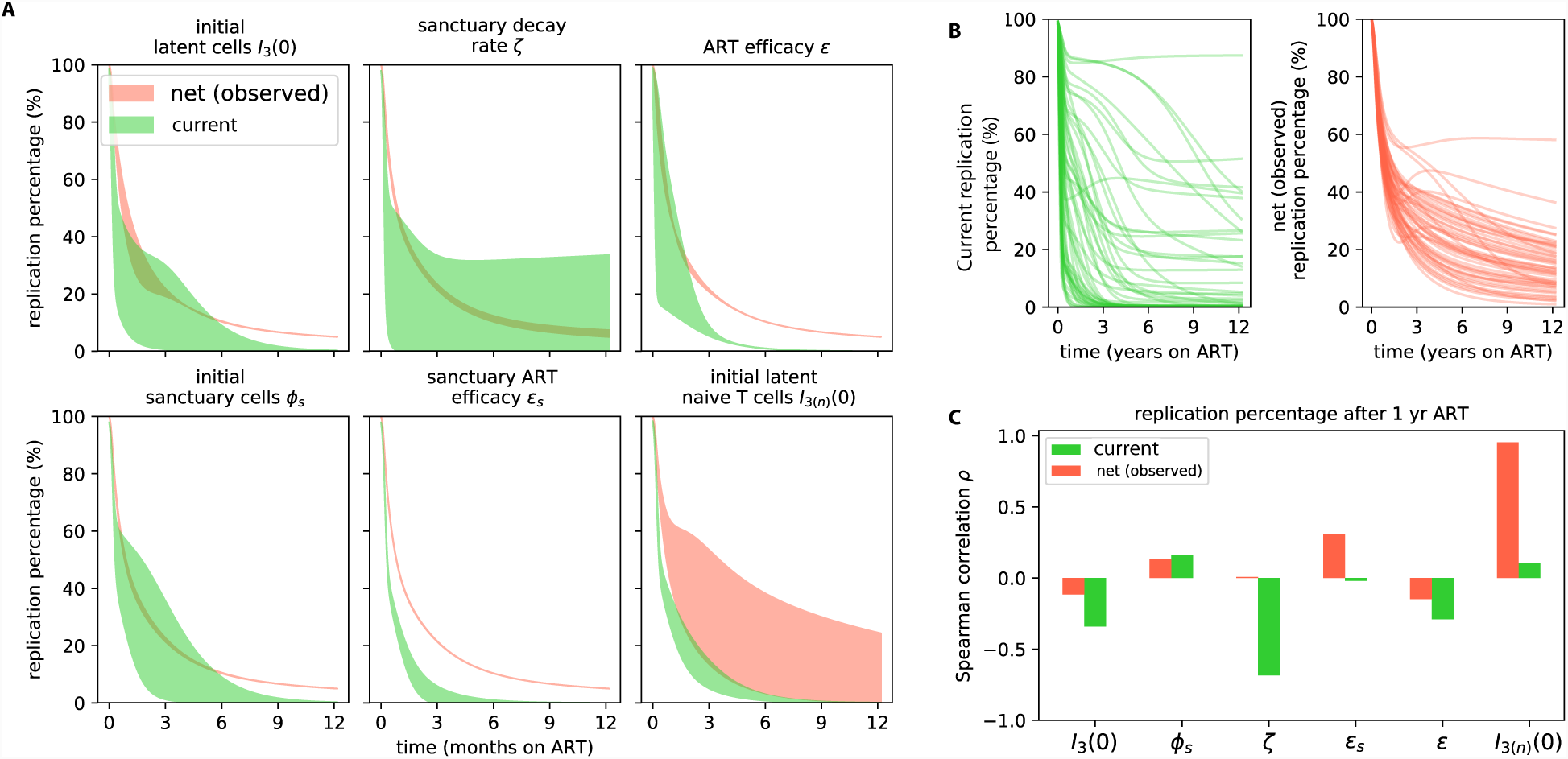
Transition from replication to proliferation as the dominant mechanism of HIV persistence during ART occurs under a wide range of parameter assumptions. **A-C**. See Methods for complete simulated parameter ranges. **A.** Local sensitivity analysis (green: current infection, red: net infection) revealed no meaningful difference in percentage of new infected cells generated by viral replication after a year of ART despite variability in initial reservoir volume I_3_ (0), sanctuary fraction φ_s_, and ART effectiveness in and out of the sanctuary (є_s_andє). Only an extremely low, or zero, sanctuary decay rate ζ predicted that a meaningful percentage (25%) of infected cells would be newly generated by HIV replication at one year, despite the fact that signals of evolution are not typically observed at this timepoint. Including a high percentage of slowly proliferating naïve CD4+ T cells (T_n_) in the reservoir alters the percentage of net, but not current, replication percentage. **B**. 50 examples from 1,000 global sensitivity analysis simulations. HIV replication accounted for fewer than 25% of current and net infected cells after a year of ART in a majority of simulations. **C.** The parameters most correlated with current and net replication percentage at 1 year of ART are different. Current replication percentage inversely correlates with sanctuary decay rate while net (observed) replication percentage positively correlates with reservoir composition (the fraction of naïve latently infected cells). Correlations are measured with a Spearman correlation coefficient.

The net replication percentage was completely unaffected by the decay rate of target cells within the drug sanctuary. Only an increase in the percentage of slowly proliferating reservoir cells (T_n_) predicted an increase in the net replication percentage (**Fig 9A**). The drivers of current infected cell and net infected cell origin therefore differed completely, highlighting the major differences between observed sequence data and contemporaneous mechanisms generating new infected cells.

To confirm these results, we simulated 10^4^ possible patients in a global sensitivity analysis in which all parameter values were simultaneously varied. A rapid transition to proliferation as the source of new infected cells occurred during year one of ART in a majority of simulated patients, and the same variables correlated significantly with net and current replication percentage, respectively (**Fig 9B&C**). Overall, this analysis does not rule out the possibility of a drug sanctuary but does confirm that its relative impact compared to cellular proliferation is likely to be minimal.

## Discussion

To eliminate HIV infected cells during prolonged ART, it is necessary to understand the mechanisms by which they persist. In this paper, we used existing data and two methods – inference of HIV clone distributions and mechanistic mathematical modeling – to determine that a majority of infected cell persistence is due to cellular proliferation rather than HIV replication. These conclusions suggest strategies that enhance ART delivery to anatomic drug sanctuaries are less likely to be effective at reducing infected cell burden relative to reservoir reduction strategies. In particular, antiproliferative therapies provide an ideal response to the observed dominance of proliferation.

In the first part of the paper, we used existing data to infer the true clonal distributions within the entire reservoir of HIV sequences in infected participants on long term ART. While the raw data indicate substantial fractions of *observed* singleton sequences, when the total reservoir size is considered, these observed singletons are revealed to be predominately members of clonal populations. In fact, the HIV reservoir appears to be defined by a rank-abundance distribution of clone sizes that can be roughly approximated as a power-law relationship. This distribution implies that a small number of massive clones, and a massive number of small clones, comprise a large percentage of sequences.

A power-law distribution can be created when a heterogeneous population grows multiplicatively with a widely variable growth rate.^55^ This suggests that the distribution of clone sizes in the reservoir is likely to have a mechanistic basis. It is plausible, though unproven, that such variable growth arises from rapid bursts of CD4+ T cell proliferation due to cognate antigen recognition. HIV integration into tumor suppression genes could also account for some observed clonal dominance.^36,37^ Smaller clones may arise from homeostatic proliferation, or less frequent exposure to smaller amounts of cognate antigen.

Another consequence of our inference is that we can more precisely define the mechanism sustaining equivalent sequences observed in longitudinal samples separated by many years. While we cannot rule out cellular longevity as a cause of HIV persistence in certain cells, the observation of multiple clonal sequences could not arise from purely long-lived latently infected cells. In fact, our analysis suggests that most observed singlet sequences arise from resampling clonal populations that have undergone many rounds of proliferation.

The first analysis does not include time-dynamics in the reservoir. Consequently, in the second part of the paper we develop a mechanistic model to reconcile observations from early and late ART. This model is the first to include the three main mechanistic hypotheses for reservoir persistence: an ART sanctuary, long-lived latent cells, and proliferation of latent cells. The model recapitulates known HIV RNA decay kinetics while tracking cells that originate from ongoing replication and cellular proliferation.

The model helps to explain how a “fossil record” of evolution would be observed early during ART, whether or not a small drug sanctuary exists. The model tracks both the fraction of cells that were generated by viral replication at a given time (current replication percentage) and the fraction that were generated by viral replication at any time point but are “fossilized” in a long-lived latently infected state (net, or observed, replication percentage). The net replication percentage remains non-negligible in the first months of ART even while the current replication percentage drops rapidly. Thus, an observed sequence that was once created by viral replication (and thus might give a signal of divergence from the founder virus) can represent a historic replication event rather than current replication. Because time of detection does not correlate linearly with sequence age, inference of evolution early during ART is problematic.^20,21^ However, the fossil record is transient: within a year of effective ART, observed phylogenetic data is more likely to represent true reservoir dynamics. Our model agrees with observations reflecting a lack of contemporaneous HIV evolution after this time.^14,22-27,29,30,36,37^

Our sensitivity analysis shows that the major variable correlating with higher observed replication percentages (a larger proportion of slowly proliferating CD4+ T cells in the reservoir) is not the same variable that correlates with higher new replication percentages (a slower decrease in sanctuary size). Replication percentage correlates with the amount of ongoing evolution in viral populations. Without requiring any phylogenetic simulation, this simple model provides an explanation for evolution during the first months of ART and no observed HIV evolution in participants with a year of ART.^14,22-27,29,30,36,37^ If we assume a large drug sanctuary and do not allow it to contract as a result of target cell decline, a persistent low-level sanctuary would emerge that stabilizes at 6 months and generates ongoing evolution at later ART timepoints. Notably, this has not been observed in clinical studies.

Our modeling results inform experiments in two ways. Using rarefaction, we suggest reasonable sample sizes to verify our hypotheses experimentally (see **Supplementary Fig 4**). We demonstrate that observed values of sequence richness and clone size, are substantial underestimates. Current studies only sample the “tip of the iceberg” of the HIV reservoir. Hundreds of thousands of infected cells from a single time point would be required to capture true reservoir diversity. This sampling depth could only be feasibly achieved as part of an autopsy study.

By using dynamical modeling, we also demonstrate that the wash-out period for the fossil record of HIV replication may be up to a year post ART. Thus, we suggest that future reservoir studies are conducted after this time point to avoid observation of historic evolution rather than contemporaneous dynamics.

The work presented here carries several important caveats. Current integration site data is still uncommon and, while robust, is limited to a handful of participants in only a few studies. Modeling rank abundance curves makes a large assumption about the continuity of the data. The power law model represents but one approach, and future work should attempt to uncover why that distribution appears to provide good fit to the data. Extrapolating abundance curves has been criticized: we note that our attempt to design a simple parametric model was based on the additional information of reservoir size and our goal to define an upper limit on reservoir richness;^56^ we also emphasize that the tail of our distributions is impossible to precisely characterize with our methods. Our approach is calibrated against sequence data from blood. However, the dynamics of HIV within lymph tissue may have different distributions. While historically, blood samples have been taken as a surrogate for HIV infected cells, we cannot rule out the possibility that the drug sanctuary that does not exchange virus or infected cells with blood. This sanctuary would be unobservable until probed anatomically. It seems unlikely that such a sanctuary could be sustained because some trafficking of CD4+ T cells from other compartments seems necessary to avoid terminal target cell limitation. However, future studies should address possible one-way trafficking or local proliferation of target cells.

In conclusion, we demonstrate that the majority of HIV infected cells arise from proliferation after the first year of ART. We have also provided an explanation for incongruent observations of evolution before and after a year of ART. Because proliferation appears to be the dominant force sustaining the HIV reservoir,^34^ we suggest limiting proliferation as a prime therapeutic target.^10,11,57^

## Methods

### Rank abundance of HIV integration sites

We used an ecological framework to study the abundance of clonal HIV. To do so, we applied methods to integration site and replication competent HIV sequence data. Cellular DNA found with HIV integrated into different integration sites in the human genome were defined as distinct “clones”. The number of times a cell was found with the same integration site added to the “abundance” of that clone. By ordering (ranking) the clones from largest to smallest by abundance, we developed a rank abundance curve, a(*r*), for each participant time point. No assumptions were made about the stability or dynamics of the reservoir rank abundance over time.

In our analysis of data from Wagner *et al.*,^37^ we combine measurements taken closely in time and use the median time point as done in that published paper. In our analysis of Maldarelli *et al.*,^36^ we grouped by integration site, or nearest measured integration site when integration site was not noted. It is important to note that the methods used by Wagner *et al.* and Maldarelli *et al.* are slightly different. The ISLA method used by Wagner *et al.* is lower throughput than the next generation shotgun sequencing method used by Maldarelli *et al.* The absolute number of viruses identified by each group therefore differs. However, the percentage of observed singletons is similar between the two studies.

We manually counted the abundance of replication competent HIV sequences using phylogenetic trees in Hosmane *et al.*^34^

### Calculation of rarefaction curves

We used rarefaction curves to estimate the expected number of distinct sequences that would still be present in a subsample of k sequences from the observed data with sample size of ***N***:

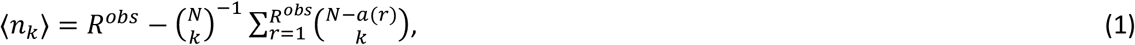

where the parentheses indicate binomial coefficients, e.g.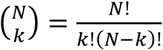. Later, we extrapolated rarefaction curves using the modeled distributions for the total reservoir size *L*. Because the number of samples we allowed was orders of magnitude smaller than the number of cells in the reservoir, *k* ≪ *L*, we used Stirling’s approximation to simplify the binomial coefficients. The expected number of sequences after k samples is then

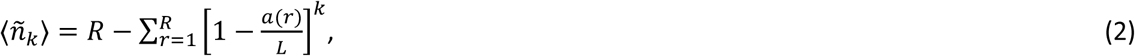

an expression which avoids computation of large factorials (derivation in the **Supplementary Methods**).

### Nonparametric estimation of species richness

We employed the Chao1 estimator to set a lower bound on the sequence or integration site richness.^58^ A derivation of the estimator is included in the **Supplementary Methods**. Chao1 is not a mechanistic model and requires no free parameters. Inference relies on only the number of observed singleton (*N*_*1*_) and observed doubleton (N_2_)sequences such that

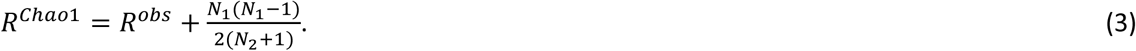

We display an asymmetric confidence interval in **Fig 3** (see Chao *et al.*^58^ or **Supplementary Methods** for the calculation). We also note it is possible the data are undersampled to the extent that a one-sided confidence interval may be more appropriate. Thus, for our biological conclusions we take the Chao1 point estimate as a lower bound, and constrain the upper bound using the parametric model (**Eq 4**). Other richness estimators (jackknife 1 and 2) were tested but provided similar and consistently lower estimates of richness than the Chao1 estimator. These were not included in our results because the Chao1 was interpreted as a lower bound on true sequence richness.

### Parametric models to extrapolate sequence abundance curves

Estimates of the size of the HIV reservoir (both replication competent and total) were gathered from the published literature.^33^ We then developed a parametric model to quantify the true rank abundance distribution of the complete HIV reservoir. Examination of the data indicated a possible log-log-linear relationship, so we chose a discrete integer power law model so that the probability of a rank is described by *p*(*r*) = *ψ*(*R*)*r*^*-α*^ where the coefficient (R) 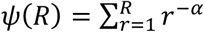is the normalization constant for the power law. Then, to describe the true rank abundance a(r) we chose the reservoir size depending on the model context (replication competent *L* = 10^7^ or total HIV DNA *L* = 10^9^). To ensure integer number of cells, we rounded this distribution, and forced the total number of cells to equal the reservoir size. That is,

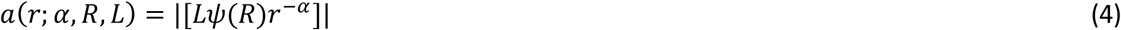

where |[]| indicates rounding to the nearest integer. Thus, our model depended on two free parameters, a power law exponent *α*, and the reservoir richness *R*. Other functional forms were explored but simplicity and accurate reproduction of the data were optimal with the power law.

### Fitting the rank-abundance model to experimental data

Using the experimental data we found the best-fit model using the following procedure. We fixed the reservoir size *L* depending on the model context (replication competent or total HIV DNA). We chose a value for *R* and α from ranges *R* ∈ [10^3^, 10^7^] and *α* ∈ [0,2] to specify the model. Then, we sampled the extrapolated distribution 10 times using multinomial sampling with the same number of samples as the experimental data being fit, *M*(*N*^*obs*^, *p*(*r*)). This procedure assumes that sampling cells does not change the distribution of the reservoir, which is reasonable given the reservoir size. Each sampled data set was compared to the experimental data by computing the residual sum of squares (rss) error of the cumulative proportional abundance (cpa) curves. For each model then, the reported error is the average rss over the 10 resamplings. Because the rss error is not symmetric across the domain of the cpa, this approach becomes similar to minimizing the Kolmogorov-Smirnov (KS) statistic: the maximum deviation between two cumulative distributions. For each experimental data set, 2500 model parameter sets were generated, and fitting results are visualized as heat maps (see **Figs 4A, 5A** for example). Because the procedure becomes computationally expensive as R > 10^7^, we did not explore values above this threshold. In theory, it is possible to have a distribution with all clones having a single member *R* = *L*, *α* = 0. For the total DNA reservoir, this value would result in R = 10^9^. However, this model was never optimal. In fact, as richness increased beyond R ≈ 10^6^, the model was no longer sensitive to *R*. Thus, it appeared that finding the best fit α was sufficient to specify the model if proper bounds on richness were included.

We excluded models where *R* < *R*^*cha01*^, but we also sought to identify an upper bound for *R*. Indeed, certain model parameter combinations are mathematically impossible. For example, for a given power law exponent, the richness is constrained below a certain value for a given reservoir size. This observation has been considered previously in ecology under the terminology of ‘feasible sets’.^59^ To determine the largest possible richness that still has the best fit, we chose the roughly constant value of α that emerged when R was large enough to be unidentifiable. Then, we noted that for large *R* it is a. reasonable approximation to allow 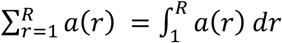. *R* is thus approximately bounded, and we solved for the maximal value or the upper bound on the richness given the best fit α and the chosen *L*. A discussion and numerical validation of this approximation is presented in the **Supplementary Methods** and **Supplementary Fig 2**. The upper bound provides the sequence abundance most permissive of true singleton sequences – the reservoir with the most evidence of HIV replication as opposed to proliferation. In extrapolated reservoirs, we used the maximum richness model to ensure we were biasing the results as strongly as possible against our own hypothesis.

### Model fitting validation with simulated data

A discussion and demonstration of model validation is included in the **Supplementary Methods** and **Supplementary Fig 1**. The exercise shows that simply fitting a power law to the experimental data (using log-log-linear regression) without the extra sampling step necessarily underestimates the power-law exponent, demonstrating the utility of our approach. Moreover, it shows that a published maximum likelihood approach^60^ is not as accurate for these data as our resampling approach (code hosted at http:11tuvalu.santafe.edu1~aaronc1powerlaws1 last accessed July 2018). We simulated a reservoir with known power law exponent **Supplementary Fig 1A** and tested for recovery of this known value. The fitting validation proceeded identically to the data fitting, 2500 distributions were generated (225 examples are shown in **Supplementary Fig 1D**), the simulated data was sampled **Supplementary Fig 1B**, and reranked **Supplementary Fig 1C**. Fitting results **Supplementary Fig 1E&F** are shown analogous to **Figs 4&5,A&B**. Finally, the most correct parameter estimation of three methods tried came from our modeling approach **Supplementary Fig 1G**.

### Mechanistic model for the persistence of the HIV reservoir

The canonical model for HIV dynamics describes the time-evolution of the concentrations of susceptible *S* and infected *I* CD4+ *T* cells and HIV virus *V*.^50,54,61^ Our model grows from the canonical model, simplifying with several approximations and extending the biological detail to simulate HIV dynamics on ART, including a long-lived latent reservoir and a potential drug sanctuary. Perelson *et al.* first noticed and quantified a ‘biphasic’ clearance of HIV virus upon initiation of ART and showed that viral half-lives of 1.5 and 14 days correspond with the half-lives of two infected cell compartments.^50,54^ With longer observation times and single-copy viral assays, Palmer *et al.* found four-phases of viral clearance after initiation of ART.^51^ Because of uncertainty in distinguishing the third and fourth phase in that study, we focus on the first three decay rates and corresponding cellular compartments, attributing a mixture of the third and fourth phase decay to the clearance of the productively infectious latent reservoir (half-life 44 months) as measured by Siliciano *et al.* and recently corroborated by Crooks *et al*.^2,3^ and the clearance of HIV DNA.^47^ We developed a mechanistic mathematical model that has three types of infected cells *I*_1_, *I*_*2,*_*I*_*3*_that are meant to simulate productively infected cells, pre-integration infected cells, and latently infected cells, respectively. We classify rapid death δ_1_ and viral production within actively infected cells *I*_1_. Cells with longer half-life that may represent pre-integration infected cells *I*_*2*_ are activated to *I*_1_ at rate ξ_2_*I*_*2*_may represent CD4+ T cells with a prolonged pre-integration phase, but their precise biology does not affect model outcomes.^48^

The state *I*_*3* (*j*)_ represents latently infected reservoir cells of phenotype *j*, which contain a single chromosomally integrated HIV DNA provirus.^44^ *I*_*3*_ reactivates to *I*_1_ at rate ξ_3_ which at present is assumed to be constant across cell phenotypes.^49^ The probabilities of a newly infected cell entering *I*_1_, *I*_*2*_,*I*_*3* (*j*)_, are *τ*_1_, *τ*_2_, *τ*_3 (*j*)_. Because we are focused on the role of proliferation, we assume sub-populations of *I*_*3*_,^12^ including effector memory (T_em_), central memory (T_cm_), and naïve (T_n_) CD4+ T cells, which proliferate and die at different rates *α*_3 (*j*)_, *δ*_3 (*j*)_. ^12,42,43^ Parameter values and initial conditions for the model are collected in **Table 1**.

### Including a decreasing sanctuary in the model

A recent hypothesis about reservoir persistence suggests there may be a small, anatomic sanctuary (1 in 10^5^ infected cells) in which ART is not therapeutic.^4^ Thus, we included the state variable *I*_*s*_ that is maintained at a constant set-point level prior to ART, where all new infected cells arise from ongoing replication. We opted for this simplification because it biased against our conclusions. The amount of virus produced by the sanctuary *V*_*s*_ is extremely low relative to non-sanctuary regions because ART results in levels undetectable by sensitive assays.^51^

Many studies have demonstrated that HIV accelerates immunosenescene through abnormal activation of CD4+ T cells.^62-64^ ART results in a marked reduction of T cell activation and apoptosis, a potential signature of HIV susceptible cells.^65^ By examining the decline of activation markers for CD4+ T cells, we approximated the decay kinetics of activated T cells upon ART, inferring approximate decay kinetics of the target cells in our model.^52,53,66^ A range of initial values exists (from ~5-20% activation) depending on stage of HIV infection, yet after a year of ART, a large percentage of patients return to almost normal, or slightly elevated CD4+ T cell activation levels (2-3%).^52^ Because we assume that target cell depletion is minimal at viral load set-point, we can approximate that the susceptible cell concentration decreases over time as the immune activation decreases, i.e., *S* = *S*(0)*e*^*-ζ t*^. This single exponential decay is simplified (it may be biphasic but the data are not granular enough to discriminate this dynamic subtlety). From existing data, the decay constant should be in the range *ζ~*[0.002, 0.01] day^-1.52,66^ We extend this decay into the sanctuary, allowing the number of susceptible cells over the whole body to decrease so that we have *I*_*S*_ = *I*_1_(0)*φ*_*S*_*e*^−*ζt*^ where *φ*_s_ is the fraction of infected cells that are in a sanctuary. Model simulations are also performed without this assumption of target cell contraction.

Last, we use the quasi-static approximation that virus is proportional to the number of actively infected cells in all compartments *V* = *n*(*I*_1_+ *Is*) where *n*= π/γ, the ratio of the viral production rate to the viral clearance rate (**Table 1**). The model is thus

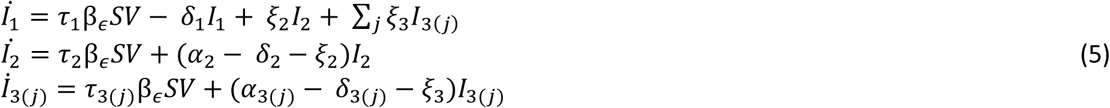

where we use the over-dot to denote the time derivative.

### Comparing proliferation and viral replication: ‘net’ and ‘current’ percentages

By solving the ODE model (**Eq 6**), we have the time solution for each infected cell state. From these, we can compute the total number of newly infected cells generated in a given time interval Δt by ongoing replication. That value is 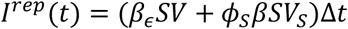The total number of newly infected cells generated by proliferation of a previously infected cell can be computed similarly in a time interval as *I*^pro^ (t) = ∑_*i* (*j*)_ *α*_*i* (*j*)_*I*_*i* (*j*)_ Δ*t*. Therefore, the percentage of infected cells generated by current replication is written

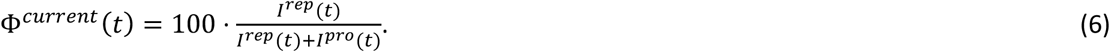

We can further subset this newly generated fraction by examining the percentage of newly infected cells that enter the long-lived latent state *I*_*3*_ by defining *I*.^*rep*(3)^(*t*)=..*τ*_3_(*β*_*∈*_.*SV* + *ϕ*_*s*_*βSV*_*s*_)Δ*t* and *I*^*pro(3)*^(*t*) = Σ_*j*_*α*_*3(j)*_*I*_*3(j)*_ Δ*t* so that

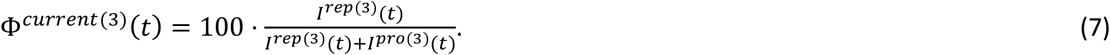

The net (or observed) replication percentage, is the fraction of cells that remain that were once generated by viral replication. To compute this quantity, we use an additional set of ODEs that we refer to as “tracking equations” because they do not change the dynamics of the system, and only are used to track specific variables. To denote the net value as opposed to new value we use a subscript Σ. The net cells generated by viral replication in state i of phenotype *j* is governed by the differential equation

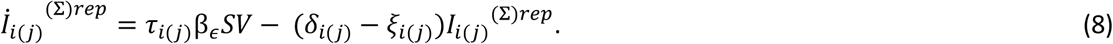

Likewise, the net cells generated by proliferation in state i of phenotype j is governed by the differential equation

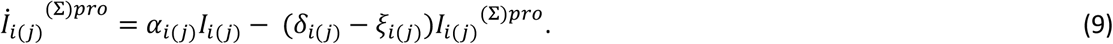

We note that because we only allow these two mechanisms, 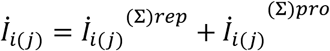 and 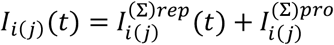. By solving the tracking equations separately, we can then find the net replication percentage by summing over cell types and phenotypes to ultimately write

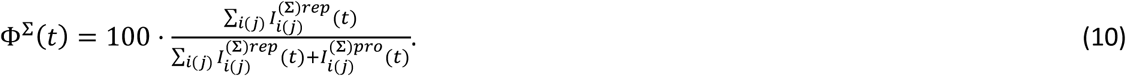

In all simulations, we assumed that 100% of infected cells at the initiation of ART were generated by viral replication, that is Φ^∑^(0) = 100. This assumption biases results in favor of replication. However, we choose it because, to the best of our knowledge, studies of proliferation during chronic untreated HIV have not been performed.

### Sensitivity analysis

Using estimated parameter bounds [lower, upper], we completed a local and global sensitivity analysis. These ranges were chosen to cover a wide range of possible assumptions. We allowed *I*_3_ (0) = [0.02,2] cells μL^-1^, *φ*_*s*_ = [10^-6^, 10^-4^] unitless, *ζ*= [0,0.2] day^-1^,*є* = [0.9,0.99] unitless, *є*_*s*_ = [0,0.9] unitless, *I*_*3* (*n*)_(0) = [0,0.5] × I_*3*_(0) cells μL^-1^. For the local analysis, we used all values as in **Table 1** and modified one parameter at a time over each listed range above. The global analysis was performed by using 10^4^ Latin Hypercube samplings of the complete 6-dimensional parameter space.^67^ The key outcome, the replication percentage (net and current) at 1 year of ART, was correlated to each parameter using the Spearman correlation coefficient—defined by the ratio of the covariance between the outcome and the variable divided by the standard deviations of each when the variables were rank-ordered by value.

### Data and code availability

Computational code for all calculations and simulations was performed in Python and Matlab and can be found at **https:11github.com1dbrvs1reservoir_persistence**. Sequence data was obtained from the Retrovirus Integration Database (RID).^68^

## End Notes

The authors declare no competing interests.

### Acknowledgments

DBR thanks F Boshier and O Hyrien for many illuminating conversations and JM Drake for originally suggesting the Chao1 estimator. DBR is supported by a Washington Research Foundation (WRF) Postdoctoral Fellowship. ERD is supported by the National Center for Advancing Translational Sciences of the NIH under Award Number KL2 TR002317. JTS is supported by NIH grants: P01 AI030371-24, U19 AI113173-02, R01 AI121129-01, UM1 AI126623, UM1 AI068635.

## Author contributions statement

DBR, ERD, and JTS conceived the study. TAW, SEP, and AMS contributed ideas and data sources for the project. DBR assembled data, wrote all code, performed all calculations, ran the models, and analyzed output data. JTS and DBR wrote the manuscript with contributions from all other authors.

